# PECLIDES Neuro A Personalisable Clinical Decision Support System for Neurological Diseases

**DOI:** 10.1101/708818

**Authors:** Tamara Müller, Pietro Lio’

## Abstract

Neurodegenerative diseases such as Alzheimer’s and Parkinson’s impact millions of people worldwide. Early diagnosis has proven to greatly increase the chances of slowing down the diseases’ progression. Correct diagnosis often relies on the analysis of large amounts of patient data, and thus lends itself well to support from machine learning algorithms, which are able to learn from past diagnosis and see clearly through the complex interactions of a patient’s symptoms. Unfortunately, many contemporary machine learning techniques fail to reveal details about how they reach their conclusions, a property considered fundamental when providing a diagnosis. This is one reason why we introduce our Personalisable Clinical Decision Support System *PECLIDES* that provides a clear insight into the decision making process on top of the diagnosis. Our algorithm enriches the fundamental work of Masheyekhi and Gras in data integration, personal medicine, usability, visualisation and interactivity.

Our decision support system is an operation of translational medicine. It is based on random forests, is personalisable and allows a clear insight into the decision making process. A well-structured rule set is created and every rule of the decision making process can be observed by the user (physician). Furthermore, the user has an impact on the creation of the final rule set and the algorithm allows the comparison of different diseases as well as regional differences in the same disease^1^.

## I. INTRODUCTION

The average life expectancy of Europeans increased by 2.9 years in the last decade. People reached an average age of 80.6 in 2013 [1] and there is more than a 50% probability that by 2030, national female life expectancy will break the 90 year barrier [2]. But a longer life does not implicate a healthy one. With higher age comes an increased likelihood of chronic diseases. This trend affects the well-being of elderly people and bears huge challenges for society and economics [1]. Computer algorithms and technology can support disease detections for example and it is hoped that systems like the one presented in this work will become increasingly prevalent as we continue to improve the state-of-the art in predictive medicine.

### A. Neurological Diseases

Alzheimer’s and Parkinson’s Disease are two of the most common neurodegenerative diseases [3][4]. In the US there are currently about 5.5 million patients of Alzheimer’s Disease (AD) and predictions project this number to grow to about 13.8 million by mid-century. In 2014 official death certificates recorded AD to be the sixth leading cause of death in the US. The average per-person medical payments for services to Alzheimer’s patients (or patients with other dementia) older than 65, are three times greater than payments for beneficiaries without these conditions [5]. The structure of Alzheimer patients’ brains changes with the disease. A larger amount of so called plaques and tangles are built by certain proteins, which lead to a loss of connections between nerve cells. This results in the death of nerve cells and a reduced amount of brain tissue. Furthermore, message transmission is less effective, as certain chemicals are missing in the patients’ brains [3][6]. Studies have shown that age is the most significant risk factor for AD [7]. But there are also genetic factors that can play a role. The Apolipoprotein E (ApoE) gene, or more specifically one of its three major isoforms, for example is known to be associated with the development of AD [8][9][10]. Different alleles of the gene can indicate higher or lower risk for developing the disease [11].

It is estimated that about 1-2% of the world population suffer from Parkinson’s Disease (PD). Almost half of the patients develop PD during age 50 and 60 [12]. A characteristic of PD is the progressive loss of substantia nigra dopamine neurons and striatal projections [10]. Consequently, patients have a lack of dopamine in their brain. The exact reason for this is mostly unknown. A consequence of dopamine shortage is that movements become slower and more difficult. Tremor, muscle stiffness, and slowness of movement are the three main symptoms of PD. Other symptoms are tiredness, pain, depression and constipation. Estimated numbers of people diagnosed with Parkinson’s in 2018 in the UK are around 145.000. Currently there is no cure for Parkinson’s disease, but treatments can control the symptoms to a certain amount. Drugs, deep brain stimulations, and physical therapies are the most common treatments [4].

### B. Decision Making

Making the right decision is one of the key factors of successfully achieving goals in all areas of work and there are numerous ways of finding the right decision. Nevertheless, the basic idea is mostly the same. It is usually a combination of experiences, research results and personal judgement. As the first two components are constantly and rapidly growing, one can imagine that decision making in general has a great potential to improve over time. But this growth also results in unmanageable amounts of data, which is why we need support systems to help processing them [13]. The goal of our support system *PECLIDES Neuro* is to integrate all three mentioned components: experience, research results and personal judgement. Especially the last aspect is rarely represented in machine learning techniques but plays an important role in every decision-making process and should therefore also be considered within a decision support system.

The following sections introduce a decision support system and cover related work, the design of the algorithm as well as the evaluation on selected data sets. Figure 1 shows an overview of our support system. After pre-processing of the data, a random forest is trained. Subsequently, a rule set is extracted from the latter and then reduced in several steps to get a smaller rule set.

**Fig. 1:**
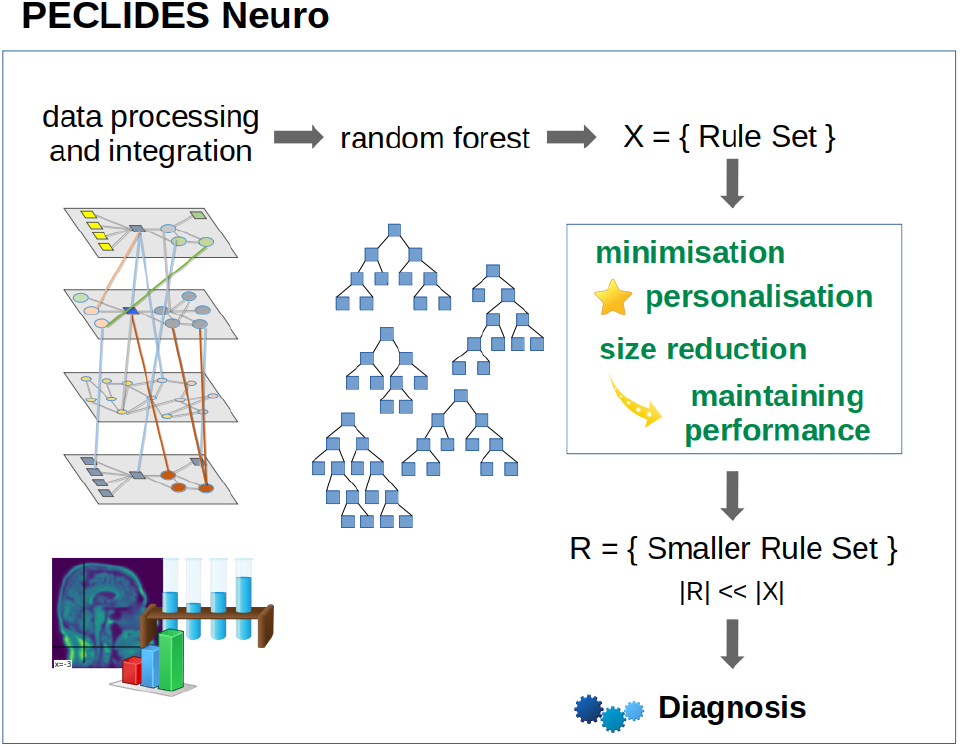
A visual overview of our clinical decision support system; data pre-processing includes feature extraction; Afterwards, a random forest is trained and converted into a rule set. This rule set is then reduced with the goal of keeping the performance and considering personal preferences in form of favourite features and enables to make a diagnosis.

## II. Related Work

Supporting medical decisions with current technology is highly discussed in literature. A frequently used technique is the ensemble learning method of random forests. Random forests are a combination of decision tree predictors and a regression and classification method. Each tree individually votes for the most popular class and their creation depends on the values of a random vector which is sampled independently but with the same distribution for all trees in the forest. In general, random forests are robust against over-fitting, run efficiently on large data and handle heterogeneous data well [14][15]. As random forests are based on decision trees, they can be used to explicitly and understandably describe a decision-making process.

### A. Rule Extraction from Random Forest

One disadvantage of random forests is that they can grow very big and become unclear. But by extracting rules from a built forest, one can gain an insight into how decisionmaking process. Mashayekhi and Gras [16] introduced two methods called RF+HC and RF+HC_CMPR which allow to extract a rule set from random forests. The main idea is to reduce the number of rules radically and therefore increase the comprehensibility of the underlying model. The rule extraction can be seen as an optimisation problem and finding the best rule set is an NP-hard problem [16].

Their proposed algorithm consists of four steps. The first one is to generate the random forest and extract all rules into a rule set. Secondly, a score for all rules is defined. For the RF+HC method they used equation 1. Hereby, *cc* stands for correct classification and is the number of covered training samples that are classified correctly. The variable *ic* refers to the incorrect classification, so the number of incorrectly classified training samples, and *k* is a predefined positive constant value. Mashayekhi and Gras proposed to set *k* = 4:

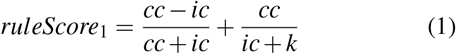

Their score leads to the elimination of noisy rules and the maintenance of rules with higher accuracy. In the third step of the algorithm a final rule set is generated. The probability of selecting a rule is hereby proportional to its score. The last step is to apply the extracted rule set on the test data set to evaluate its performance. The average rule set size after applying the RF+HC algorithm is 0.6% of the original rule set, while the accuracy generally only decreases by a couple of percent [16].

Their second method, called RF+HC_CMPR, is an extension of the firstly proposed method RF+HC, where they additionally considered the length of the rules. It adds another addend to *ruleScore*_1_. Equation 2 shows the new score:

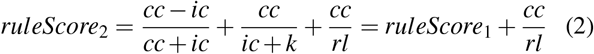

Hereby *rl* refers to the length of the rule. This way a shorter rule adds a higher number to the original score than a long rule, which leads to a higher score. The purpose is to favour shorter rules, as they are more transparent and understandable than longer rules. Mashayekhi and Gras applied their two algorithms to several different data sets and compared them to CRF and the “normal” Random Forest methods. CRF is a method that was introduced by Liu et al. It is a joint rule extraction and feature selection method [17] [18]. On average, both the RF+HC and the RF+HC_CMPR method resulted in almost the same accuracy as the CRF method. Furthermore, on average over all data sets, the three methods obtained 96% of the random forest accuracy. Moreover RF+HC and RF+HC_CMPR resulted in a clearly smaller number of rules. On average, the size of the extracted rule set is 0.6% of the original random forest. In comparison, the total number of rules in CRF is 11.66% of the size of the original rule set [16].

### B. Shrinking Random Forests

Even though decision trees represent a fairly transparent way to make decisions, random forests can get very big and quite incomprehensible. Therefore it is useful to consider different ways of shrinking a random forest while maintaining its prediction accuracy. To minimise the size of a random forest, one has to decide when and which trees can be eliminated. Zhang and Wang [19] presented three different measures to determine the importance of a tree. (1) A tree is not necessary, if its removal from the forest has the least impact on the overall prediction accuracy. Furthermore, a tree can be removed, if it is highly similar to other trees in the forest. This similarity can be either measured as (2) an average similarity to all other trees or (3) a pairwise similarity.

To get the tree with the least impact on the forest (1), firstly the prediction *p_F_* of the whole forest is calculated. Secondly, for each tree *T* in the forest *F* the prediction *F_−T_* of the forest without the tree *T* is determined. Lastly, the tree that leads to the smallest difference in prediction accuracy

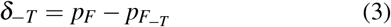

can be removed.

The similarity between two trees can be defined by the correlation between their predicted outcomes. The average similarity (2) can be calculated as following:

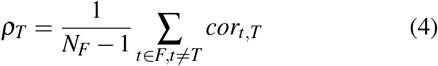

where *N_F_* is the number of trees in the forest and *cor_t,T_* is the correlation between two trees *t* and *T*. The tree T with the highest *ρ_T_* has the highest similarity to the rest of the forest and can therefore be eliminated.

The highest pairwise similarity (3) is measured by the correlation of the accuracy of two trees. Firstly, a weight *w_T_* is introduced for every tree T and set to 1. Subsequently, one is searching for the two trees *T*_*s*1_ and *T*_*s*2_, which are most similar. Afterwards, the average of similarity *ρ*_*s*1_ and *ρ*_*s*2_ for those two trees is calculated. The tree *T^rs^* with higher *ρ* can then be removed. Finally, the following weights are calculated:

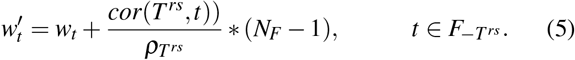

As a last step, it is important to select the optimal size for the sub-forest. Zhang and Wang proposed to define a performance trajectory *h*(*i*),*i* = 1,…,*N_f_* − 1 of a sub-forest of *i* trees, where *N_F_* is the size of the original random forest. The optimal size can then be selected by maximising *h*(*i*) over *i* = 1,…,*N_F_* − 1 [19].

Zhang and Wang showed on real data sets, that a shrunken random forest can sometimes even outperform the original one and oftentimes achieves a very similar accuracy to the original random forest [19].

## III. PECLIDES Neuro

Our implemented clinical decision support system is based on the machine learning technique of random forests. The first step of the algorithm is to generate such a random forest from a given data set and subsequently extract rules from it. The usage of the *RandomForestClassifier* Class in Python allows the variation of several parameters. In this project, we focused on the calibration of the number of trees in the forest, the function to measure the quality of a split, the maximum number of features to consider when looking for the best split, and the maximum depth of a tree.

To start the training process of the random forest, the data set was separated into a training and a test set. One approach we used is to generate a train-test split which takes a certain percent of the whole data set (for example 30%) aside for the test set and trains the model on the remaining samples (70%). Using 10-fold cross validation, the performance on the data set can be measured while reducing the risk of over-fitting.

### A. Building Rule Set from Random Forest

The next step after the creation of the random forest, is to extract rules from the latter. These rules will then build the core of the decision support system. This can be done by iterating through each tree in the forest and extracting each branch. Figure 2 shows an exemplary decision tree. One branch represents one rule, as it determines the decision process from the tree’s root to one leaf. Thus, the displayed tree would lead to eight rules, since it has eight leaves. In each node of the tree, a question is asked like “If *feature_X_* is smaller than Y, then left, else right”. Therefore, for each node the following three values are important: (1) the *feature_X_* that is considered, (2) the value Y that it is checked against and (3) whether the feature has to be smaller or greater than that value. So each node can be represented as a list of three elements [feature, l/g, value], whereby l/g determines whether the feature has to be lower or greater than the value. Finally, the outcome of a branch has to be stored too. This can be done by adding another element to the rule list. This outcome can be the decision whether a person has a certain disease or not. In my case the trees’ leaves determine whether a person is predicted to have Alzheimer’s or Parkinson’s Disease or whether the person is predicted to be healthy. All rules of all branches of all trees can then be stored in a set that represents the whole rule set.

**Fig. 2:**
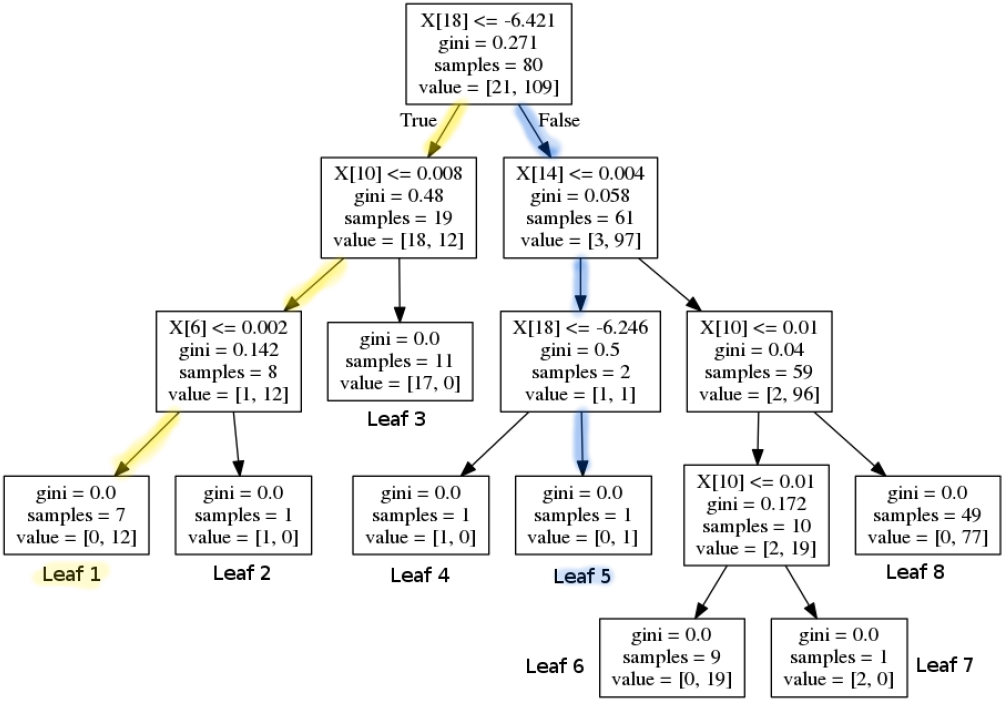
A sample decision tree that would lead to eight rules; the yellow and blue path show two exemplary rules that would be extracted from the decision tree.

Each branch can be represented by a disjunctive normal form (DNF) where all queries of all nodes are connected with ’and’. So the exemplary rule from above can be represented by the following DNF:

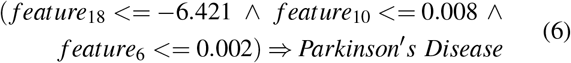

### B. Minimising the Rule Set

As the number of trees and their depths determine the number and the length of the rules, the exact size of the rule set varies from application to application. But in general, the goal is to keep the rule set small. Figure 2 shows an exemplary decision tree. One can see that feature 18 is checked twice in the blue ’False/True’-branch that leads to leaf five. The matching DNF is shown in equation 7. In the tree’s root it is checked whether feature 18 is greater than −6.421 and two layers below it is checked whether feature 18 is greater than −6.246:

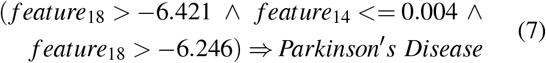

So the first logical step to minimise the rules is to eliminate redundant queries within a rule, like in our example branch from above. Deleting the first query (whether feature 18 is greater than −6.421) does not change the outcome of the rule. The condition that feature 18 has to be greater than −6.246 already implies that condition *feature*_18_ > −6.421 is fulfilled. So the mentioned branch (see equation 7) can be reduced to the following logical statement:

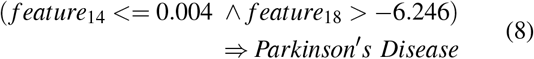

### C. Personalisable Rule Score

As suggested by Mashayekhi and Gras [16], we also assigned a score to each rule, depending on its performance on the training set. The score chosen in this work is an extension of *ruleScore*_2_ introduced in their paper (see equation 2). It considers correct and incorrect classifications as well as the length of a rule and a personalisable attribute. This score is then calculated for every rule separately. The consideration of a personal preference for features enables the rule score to be personalisable and therefore to adjust and influence the support system after the user’s preferences, experience, and expertise. The physician can nominate a desired number of features that shall be preferred during the minimisation steps and kept in the rule set. These features could be seen as extremely relevant for predicting whether a patient is diseased or not. The personalisation can lead to a more suitable support system for physicians and therefore increase the trust in the system and maybe also the willingness to use it in the first place.

Therefore *ruleScore*_2_, which was proposed by Mashayekhi and Gras [16], was extended by another component. This additional element is the consideration of one or more preferred features. If a rule does not contain a nominated favourite feature, the score will be the same as rule *ruleScore*_2_ (see equation 2). But if the rule contains a favourite feature, another value is added to increase the score. This way rules that should be preferred because they contain a favourite feature, get a higher score than others, even if the performance is the same. And as rules with a low score are eliminated first, this leads to the fact that rules which contain favourite features are less likely to be deleted from the rule set.

Equation 9 shows the new rule score. If several features are defined as preferred ones, than there will be a ranking among them. The features are a parameter list where the first list entry is the most preferred feature and the last list element the least preferred one. If a rule contains the most important feature, the rule gets a higher score than if it contains the second most important feature and so forth. The score will be the highest, if a rule contains all preferred features (considering the same performance). We decided to use a linear function to calculate the additional score points. The equation to calculate the new rule score is as following:

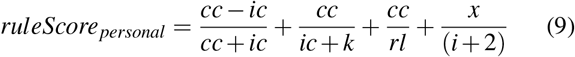

where *i* refers to the feature’s index in the list and *x* is a constant positive value that can be customised. So the first feature (index 0) increases the score by 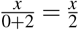, whereas the second feature (index 1) increases the rule by 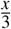 and so on. This way the favourite features are ranked and depending on how many and which of those features are considered in one rule, the rule gets a higher or lower score. The parameters *ic*, *cc*, and *k* are the same as suggested by Mashayekhi and Gras [16]. *cc* and *ic* stand for correct and incorrect classification, respectively and *k* is a positive constant value. The usage of *k* = 4 was suggested by Mashayekhi and Gras. The constant *x* can be adjusted depending on how much impact one wants the preferences to have on the whole score.

Now the rule set can be minimised to the best performing rules with the most occurrences of the preferred features. For that, one can either define a certain percentage of the rule set, that shall be deleted or kept, or a threshold to what minimum performance the rule set should be reduced. The latter holds the challenge that the performance of a rule set is not proportional to its size. It is more straightforward to reduce the rule set to a certain percentage, like 10% of the original rule set’s size and then analyse the performance of the smaller and more accessible rule set. Additionally to getting a prediction, by inspecting this small rule set, one can see which features are monitored and which thresholds are important. This can lead to information about the disease, its characteristics, and possible factors for diagnoses.

### D. Evaluation on Different Data Sets

We used a variety of data sets like Alzheimer’s Disease Neuroimaging Initiative (ADNI) [20], PROPAG-AGEING [1], a data set of spiral drawings [21] and biomedical voice measurements [22]. The data is partly a complex collection of different data types and from different institutions in Europe like PROPAG-AGEING.

The ADNI data set contains different types of data, including PET images, MR images and clinical data. Firstly, the clinical data with information about age, ethical background, gender, numerous numeric test results and more was used. All values are numerical or can be transformed into numeric values easily, which makes the implementation of a random forest straightforward. The data set consists of 3445 samples from Alzheimer’s patients and healthy subjects. The random forest achieved an accuracy of 97%, a sensitivity of 96% and a specificity of 97.4%. The rule set extracted from the random forest contains 1447 rules and the performance is strongly depending on the way the rule set is evaluated. The exhausting approach that predicts one outcome for each rule for each data sample, achieves only an accuracy of 70%. Whereas the tree-like approach, which only predicts an outcome, if the rule is applicable to the data sample, achieves an accuracy of 96.92%, which is almost as high as the performance of the random forest. The sensitivity of the tree-like approach is 96.2% and the specificity 97.24%.

After deleting the 500 weakest rules of the rule set without preferring any features, the accuracy is still very high with 97%, the sensitivity is 96.5% and the specificity 97.2%. Going down to half the rule size with 724 rules, results in an accuracy of 94.7%, a sensitivity of 86.76% and a specificity of 98.11%. Which shows again, that shrinking the rule set the right way does not have a big impact on the performance.

ADNI also provides three-dimensional *T*_1_-weighted magnetic resonance imaging (MRI) for developing and testing analysis techniques for extracting structural endpoints. To ease the utilisation of the MR Images, standardised analysis sets of data comprising scans that met minimum quality control requirements were created within ADNI. In this work, samples from 1-year completers were used, that includes images from subjects who had 6- and 12-month scans [23]. The images typically consist of 256 × 256 × 170 voxels with a voxel size of 1*mm* × 1*mm* × 1.2*mm* [24]. 27 images from Alzheimer’s patients and 25 images from healthy subjects were used.

In order to train the random forest, features have to be extracted from the MR images. Using Python and the libraries *nipy* [25] and *nilearn* [26] MR images can be processed more easily. They can be converted into arrays containing the image’s colour values. Figure 3 shows the three middle slices of an exemplary MR image (sagittal, coronal and axial plane).

**Fig. 3:**
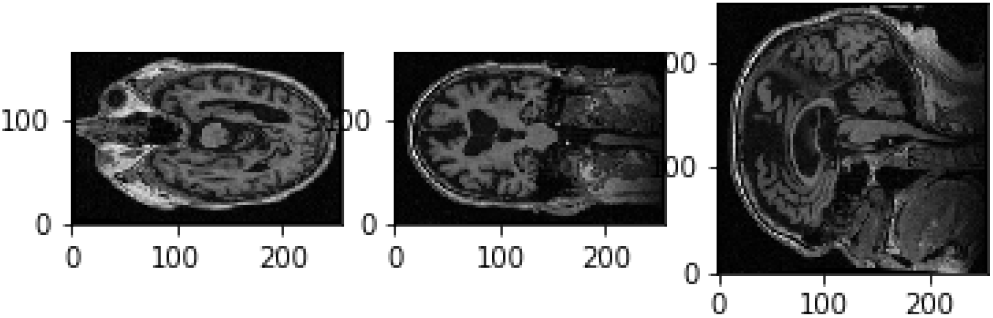
Three slices of a 3D MR image; with sagittal, coronal and axial plane

The features extracted in this work, include first order and second order descriptors [27]. Firstly, as a pre-processing step, a Gaussian filter was applied to the images (see figure 4). The filter is applied along the three first dimensions of the image [28]. The result is an array of arrays of arrays with one value for each voxel. Afterwards, the MR images are converted into arrays containing numerical values representing the colour of each voxel. Figure 4 shows three slices of a smoothed image after the application of the Gaussian filter. From these pre-processed arrays features were extracted. The first order descriptors include the sum, the mean and the maximum of all voxel values, as well as mean, sum and maximum values of the middle slices of all three dimensions. Exemplary slices are shown in figure 3. A colour level histogram was used to extract the most frequent voxel value in the MR image. The value 0 was excluded form the histogram, as the background is represented by 0 and should not be considered.

**Fig. 4:**
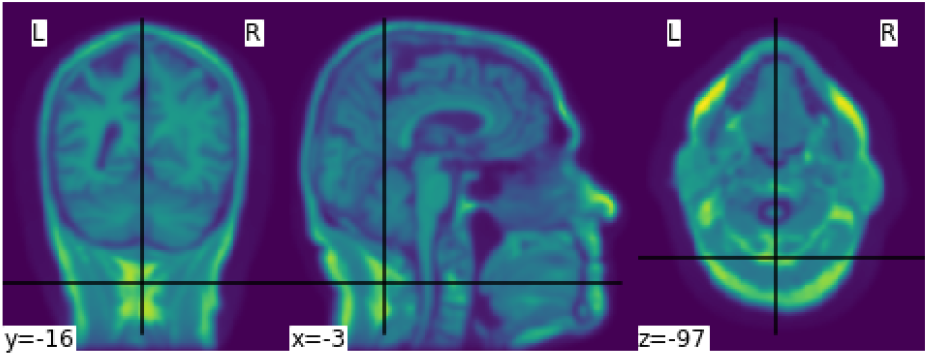
Smoothed MR image with Gaussian filter

To extract second order features, the probability of the different brain tissues were determined. The three types of tissue are cerebrospinal fluid (CSF), grey and white matter. Figure 5 shows the probabilities for the different tissue types in an exemplary slice of an MR image. The segmentation was performed using the python library *dipy* [29], its class *TissueClassifierHMRF* and the Markov Random Fields modeling approach. The latter is frequently used in literature. An example would be Held et al. who described a fully automated 3D segmentation technique for MR images [30]. The maximum a-posteriori Markov Random Field approach uses iterative conditional models and expectation maximisation to estimate the parameters [31]. After the segmentation, more features can be extracted depending on the tissue type. The sum of all voxel values separated by tissue type, as well as the maximum of the sum of the inner arrays were calculated. All features were based on the smoothed images. Using the images without this pre-processing step leads to poorer results.

**Fig. 5:**
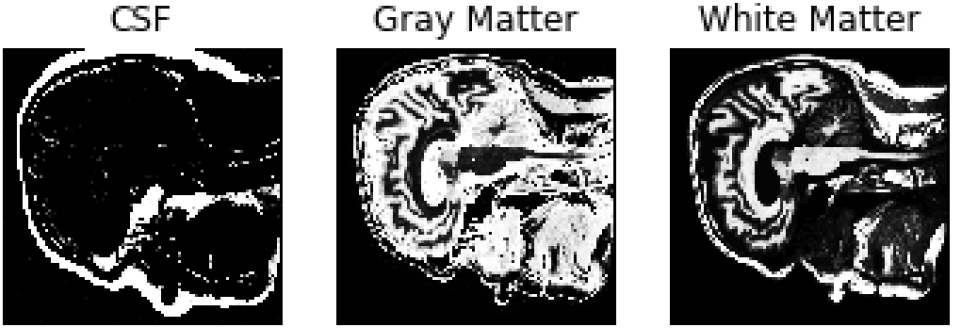
Probabilities of the different brain tissues CSF, grey matter and white matter

**Fig. 6:**
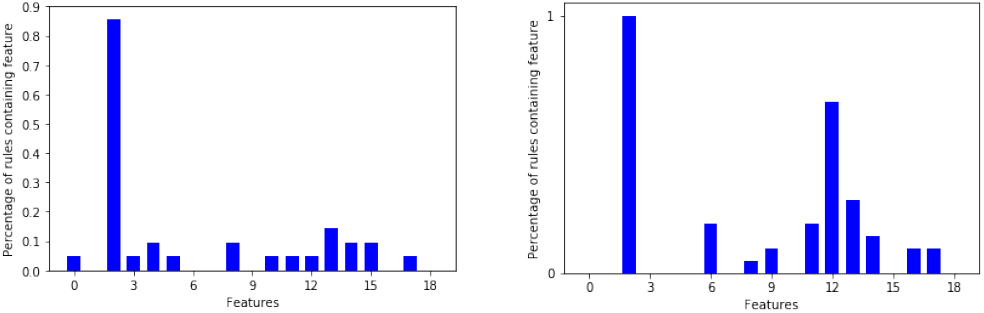
Distribution of occurrences of features in rules of MRI data set

These features were then used to implement a random forest. 10-fold cross validation lead to an average accuracy of 99.38%, a sensitivity of 99%, and a specificity of 100%. Extracting a rule set from a parameter pruned random forest, leads to a rule set of 213 rules with an accuracy, a sensitivity and a specificity of 100% each. The first reduction step that eliminates queries within singular rules (see section III-B) does not change the outcome. Table I shows some results of different rule sets with and without preferred features. Interestingly, even one rule alone can predict the correct outcome for each data sample. This feature is the maximum of the sum of the sum of the smoothed data. This feature defines a clear threshold between AD patients and the healthy control group. This might be due to the small data set and has to be tested on larger data sets again. But it most certainly is a very strong indicator to make a diagnosis about AD from MR images. Table I also shows that preferring certain features can have a greater impact on which rules are deleted than the actual performance of the rules. By reducing the rule set to 1% of its original size with three preferred features, the performance is clearly poorer than without setting a preference for features.

**TABLE I:**
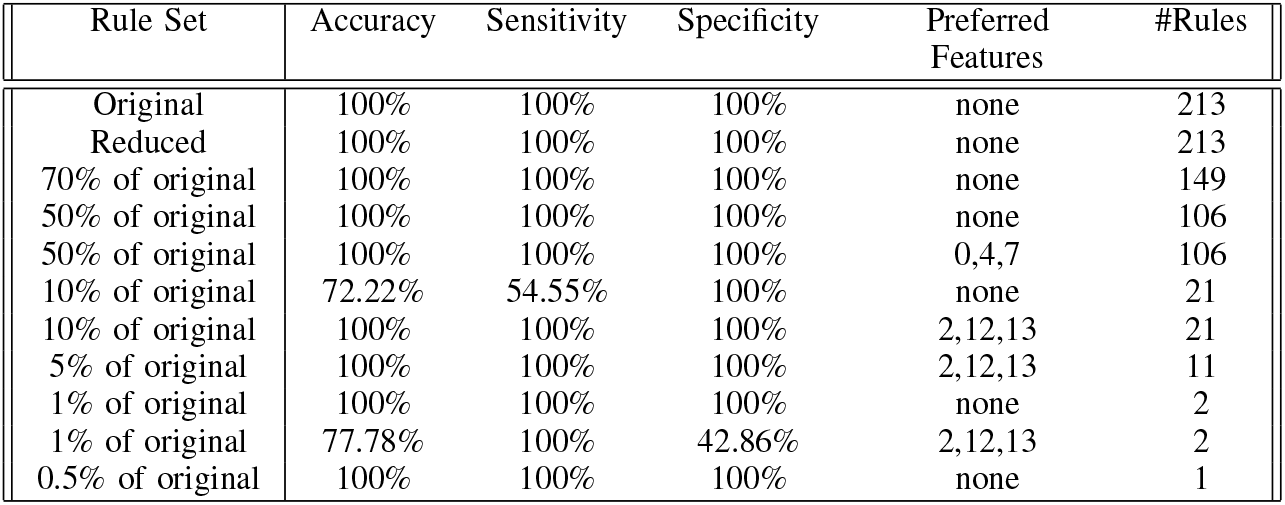
Results of different rule sets with tree-like application on MR images in ADNI data set

| Rule Set | Accuracy | Sensitivity | Specificity | Preferred Features | #Rules |
| --- | --- | --- | --- | --- | --- |
| Original | 100% | 100% | 100% | none | 213 |
| Reduced | 100% | 100% | 100% | none | 213 |
| 70% of original | 100% | 100% | 100% | none | 149 |
| 50% of original | 100% | 100% | 100% | none | 106 |
| 50% of original | 100% | 100% | 100% | 0,4,7 | 106 |
| 10% of original | 72.22% | 54.55% | 100% | none | 21 |
| 10% of original | 100% | 100% | 100% | 2,12,13 | 21 |
| 5% of original | 100% | 100% | 100% | 2,12,13 | 11 |
| 1% of original | 100% | 100% | 100% | none | 2 |
| 1% of original | 77.78% | 100% | 42.86% | 2,12,13 | 2 |
| 0.5% of original | 100% | 100% | 100% | none | 1 |

Figure 5 shows the percentage of occurrences of features in rules in the 10% rule set. For sub figure (a) no features were preferred whereas the rule set in figure (b) preferred features 2,12 and 13, the three most often represented features in the original rule set. One can see, that due to the preference of these three features, they appear more often in the shrunken rule set.

### E. Handling missing values

The rule based algorithm depends on the previous implementation of a random forest. The random forest itself does not handle missing values, but Python’s *sklearn* [32] tool provides a class called *Imputer* [33] that can handle missing values and replace them with either the mean value, the median, or the most frequent value in the respective column.

This way the support system can deal with missing values and incomplete data samples do not have to be deleted. This was for example applied in the PROPAG-AGEING data set, as some values were missing in a few data samples. One could take this into account during the reduction step of the rule set, by not choosing those less reliable features as preferred ones. This shows another advantage of the fact that the support system is personalisable.

### F. Graphical User Interface

To make the interaction with the decision support system approachable and straightforward, a user interface was implemented. Figure 7 shows the provided GUI. The first rule set that was extracted from the whole random forest, was already created beforehand. The name of the data set and the number of rules of this original rule set are stated on top of the window. Beneath, the first reduction step can be performed by clicking on the button *First Reduction*. This does not change the number of rules but eliminates redundant queries within one rule. The new size of the rule set is then stated under the button. In the next step, optionally favourite features can be named (1,2 and 3 in the example) and the percentage to which the rule set shall be reduced (here: 30%). The new size as well as accuracy, sensitivity and specificity will then be calculated and displayed after the button *Reduce Rule Set* is clicked. To make a new prediction, a value has to be filled in for each feature and with the button *Predict* a prediction is calculated and displayed (here: healthy). The two buttons *more info* open a message box with more information about what to fill in the entry boxes. The button *Print Rules* shows all rules within the current reduced rule set in form of if statements (see figure 8). The two buttons on the bottom of the GUI show bar charts with the number of rules containing each feature in the original rule set and the current reduced one (see figure 9). These two properties allow a clear insight into the decision-making process as well as the influence of different features.

**Fig. 7:**
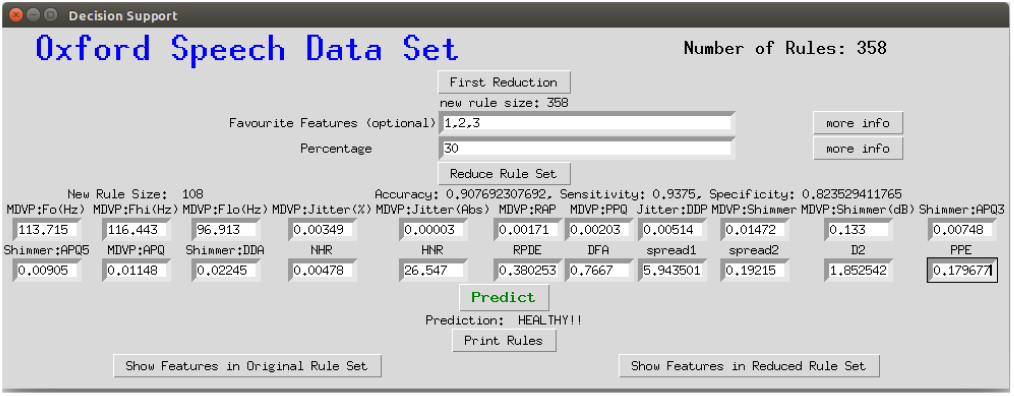
Graphical interface for the clinical decision support system

**Fig. 8:**
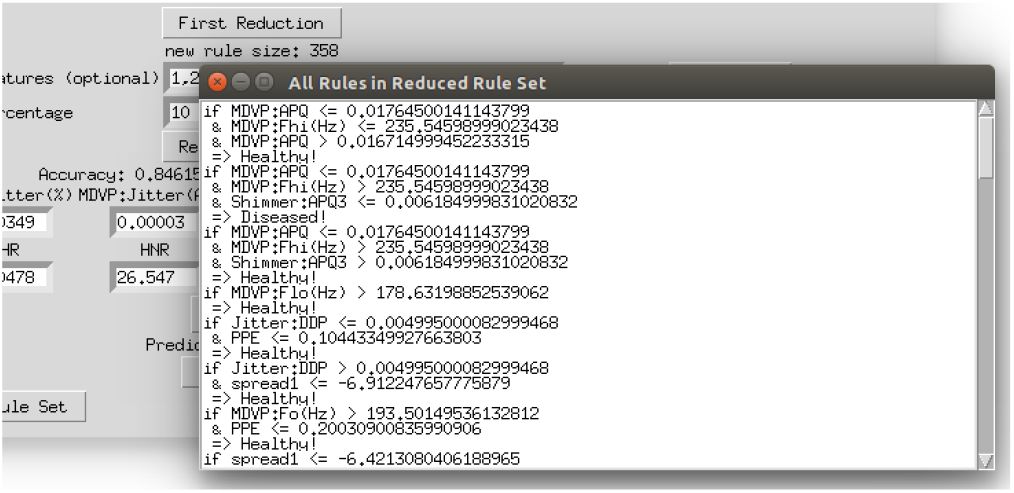
Displaying all rules that remained in the reduced rule set

**Fig. 9:**
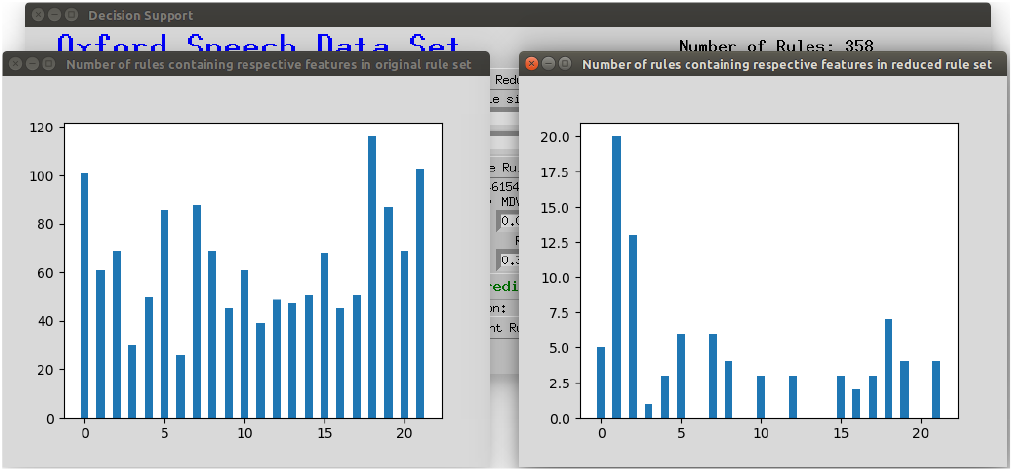
Displaying the number of rules containing the respective feature in the original rule set and the current reduced rule set

## IV. Conclusions

The algorithm introduced in this work can be used as a clinical decision support system. It helps analysing clinical data and integrating different data types and predicts the health of a subject. The focus of this work is thereby on neurological diseases like Alzheimer’s and Parkinson’s Disease. The goal is to make machine learning algorithms more transparent and accessible while ensuring a high performance. This is especially useful in clinical contexts, where a comprehensible decision-making process increases the trust in the diagnosis and can reveal information about diseases. Therefore the algorithm includes the option of personalising and adjusting the treatment of parameters within the algorithm. This leads to more insight into features used by the algorithm, their importance and informative value.

The algorithm can be divided into three major steps. (1) Firstly, a random forest is created that builds the foundation of the algorithm. (2) Secondly, a set of rules is extracted from the random forest. (3) And thirdly, this rule set is reduced using different algorithms. The third step includes the personalisable aspects, where as important considered features can be preferred within the rule set. Figure 1 shows a graphical overview of the algorithm.

The rule based algorithm allows to draw conclusions about the impact of certain features on the decision-making process. Physicians in different regions can adjust the algorithms to their patients’ needs, their experience and expertise. This might lead to different algorithms in big international cities compared to rural regions or different continents. Information about regional differences can then be used to get a better understanding of a disease in different demographic groups. Another interesting analysis could be the similarity between different diseases. Therefore knowledge about the importance of specific factors in two or more diseases are very valuable. By examining the impact of various genes for example, one could reason about the similarity between different diseases or draw conclusions about disease ontologies. Another extension of the personalisation process could be that doctors are allowed to add own rules, and adjust them to match the local aspects like ethnic groups, regional lifestyle, environment factors, or common social interactions. This would lead to a further personalisation through the interaction with individual patients.

## V. Future Work

There are several possibilities of combining random forests and neural networks, e.g. [34] [35] [36]. Those two machine learning techniques have almost complementary advantages and disadvantages. For example the knowledge representation of decision trees are mostly comprehensible whereas the decision process of neural networks is hard to understand. On the other hand, decision trees have trouble dealing with noise, which is not a big problem for neural networks. So the idea came up to combine these two approaches to benefit from both their advantages [13]. Zorman et al. introduced an idea of how to combine decision trees and neural networks. Firstly, they generated a decision tree which is then used to initialise the neural network. Subsequently, the neural network is again converted into a decision tree, which has a better performance than the original one. The resulting decision tree may not have the same performance as the neural network, but it is easier to interpret and comprehend which can be a huge benefit [37] [35] [13]. This approach could also be used for the clinical decision support system introduced in this work and is a promising approach to make decision support systems more accessible. v Random forests can also be used to initialise deep feedforward neural networks where the network’s structure is determined by the structure of the trees. These so called “deep jointly informed neural networks” (DJINN) show a warm-start to the neural network training process and result in lower cost and a lower number of user-specified hyper-parameters needed to create the neural network [36]. This shows another possibility to combine random forests with neural networks and combine both methods’ advantages. Our on random forests based decision support system could also be used as a foundation for further developments of DJINNs and bears numerous similar possibilities for extensions and on top built systems.

## Footnotes

1 The source code of Peclides Neuro can be found on GitHub: https://github.com/tamaramueller/Peclides-Neuro

